# Light-induction of endocannabinoids and activation of *Drosophila* TRPC channels

**DOI:** 10.1101/2021.06.17.448894

**Authors:** Takaaki Sokabe, Heather B. Bradshaw, Makoto Tominaga, Emma Leishman, Craig Montell

## Abstract

*Drosophila* phototransduction represents a classical model for signaling cascades that culminate with activation of TRP channels. TRP and TRPL are the canonical TRP (TRPC) channels, which are gated by light stimulation of rhodopsin and engagement of Gq and phospholipase Cβ (PLC). Despite decades of investigation, the mechanism of TRP activation in photoreceptor cells is unresolved. Here, using a combination of genetics, lipidomics and Ca^2+^ imaging, we found that light increased the levels of an abundant endocannabinoid, 2-linoleoyl glycerol (2-LG) *in vivo*. The elevation in 2-LG strictly depended on the PLC encoded by *norpA*. Moreover, this endocannabinoid upregulated TRPC-dependent Ca^2+^ influx in a heterologous expression system and in dissociated ommatidia from compound eyes. We propose that 2-LG is a physiologically relevant endocannabinoid that activates TRPC channels in photoreceptor cells.

## Introduction

Phototransduction is crucial in animals ranging from worms to humans as it enables positive or negative phototaxis, entrainment of circadian rhythms and reception of visual information from the surrounding environment. In some photoreceptor cells, such as mammalian rods and cones, the phototransduction cascade culminates with closing of cation channels (Yau & Hardie, 2009). In contrast, the cascade in *Drosophila* photoreceptor cells and in mammalian intrinsically activated retinal ganglion cells leads to opening of the cation channels (Montell, 2012; Montell, 2021; Yau & Hardie, 2009).

In *Drosophila*, the main site for light reception and transduction is the compound eye, which is comprised of ∼800 repeat units called ommatidia. A single ommatidium harbors eight photoreceptor cells, each of which includes a rhabdomere. This specialized portion of the photoreceptor cells consists of thousands of microvilli, thereby enabling rhodopsin to be expressed at very high levels for efficient photon capture (Montell, 2012; Montell, 2021; Yau & Hardie, 2009).

The *Drosophila* phototransduction cascade has been studied for over 50 years beginning with the seminal work by Pak and colleagues (Pak *et al*, 1970). This has led to the elucidation of the critical signaling proteins that transduce light into an electrical signal (Hardie & Juusola, 2015; Montell, 2021). Light-activation of rhodopsin engages a heteromeric Gq protein and stimulation of the phospholipase C (PLC) encoded by *norpA* (Bloomquist *et al*, 1988), which in turn induces opening of the Ca^2+^ permeable cation channels. These include the Transient Receptor Potential (TRP) channel, which is the classical member of the TRP family (Hardie & Minke, 1992; Montell & Rubin, 1989), and a second canonical TRP channel (TRPC), TRPL (Niemeyer *et al*, 1996;

Phillips *et al*, 1992). Related TRP channels are conserved from flies to humans (Wes *et al*, 1995; Zhu *et al*, 1995). Extensive studies of fly photoreceptor cells have also revealed mechanisms underlying the dynamic movements of signaling proteins, as well as proteins that function in the visual cycle, post-translational modification of signaling proteins, and the composition of the signalplex, which is a large macromolecular assembly that clusters together many of the key proteins that function in phototransduction through interactions with the PDZ-containing protein, INAD (Hardie & Juusola, 2015; Montell, 2012; Montell, 2021). The fly eye has also provided an outstanding tissue to model human diseases (Lin *et al*, 2018; Liu *et al*, 2017; McGurk *et al*, 2015; Pak, 1994; Warrick *et al*, 1998; Zhuang *et al*, 2016).

Stimulation of PLC is essential for activation of the TRP and TRPL channels. However, the mechanism linking activity of PLC to opening of TRP and TRPL is still under debate. PLC hydrolyzes phosphatidylinositol 4,5-bisphosphate (PIP_2_) to release inositol phosphate 1,4,5 trisphosphate (IP_3_), a H^+^ and diacylglycerol (DAG). Consistent with this conclusion, in fly heads light exposure causes a decrease in PIP_2_ levels and an increase in DAG (Huang *et al*, 2004). IP_3_ and the release of Ca^2+^ from the endoplasmic reticulum does not seem to play a role in *Drosophila* phototransduction (Acharya *et al*, 1997; Raghu *et al*, 2000). Rather, multiple other models have been proposed.

According to one study, PIP_2_ depletion accompanied by local intracellular acidification by H^+^ promotes channel activation (Huang *et al*, 2010). DAG has also been reported to activate TRP and TRPL in excised rhabdomeric membranes (Delgado & Bacigalupo, 2009; Delgado *et al*, 2019; Delgado *et al*, 2014). Mechanical contraction of the rhabdomeral membrane has been proposed to contribute to activation of the channels (Hardie & Franze, 2012). This may occur due to cleavage of the head group of PIP_2_, leaving the smaller lipid, DAG, in the membrane (Hardie & Franze, 2012).

Polyunsaturated fatty acids (PUFAs) have been reported to activate TRP and TRPL (Chyb *et al*, 1999; Delgado & Bacigalupo, 2009; Lev *et al*, 2012), although a more recent study showed that PUFAs do not increase with illumination (Delgado *et al*., 2014).

In this work, to identify physiologically relevant lipids that could activate TRP and TRPL we performed a lipidomic analysis using fly heads exposed to light or that were maintained in the dark. We found that several lipids increased in concentration upon light stimulation in control flies but not in the *norpA* mutant that eliminates the PLC required for phototransduction. The lipids that were upregulated by light included endocannabinoids and an *N*-acyl glycine (NAG). Endocannabinoids are related to plant-derived cannabinoids, which in mammals are capable of activating the same receptors as cannabinoids, such as the G-protein coupled receptors, CB1 and CB2 (Gregus & Buczynski, 2020). However, *Drosophila* has no CB1 and CB2 homologs (McPartland *et al*, 2001), and no cannabinoid receptor has been identified in flies. We found that the endocannabinoids and the NAG activated TRPL channels *in vitro* in a dose-dependent manner and induced Ca^2+^ influx in dissociated ommatidia via the TRP and TRPL channels. One endocannabinoid, 2-linoleoyl glycerol, was ∼60—100 times more abundant than the other lipids that increased upon light stimulation. We propose that 2-linoleoyl glycerol is a key lipid that contributes to activation of the TRPC channels in photoreceptor cells.

## Results

### Endocannabinoids and NAG are upregulated by light

To evaluate lipids that are increased by light stimulation of *Drosophila* photoreceptor cells, we performed lipidomic analysis (Fig 1A). In addition to using control flies (*w*^*1118*^) maintained in the dark or stimulated with light, we also analyzed *norpA*^*P24*^ mutant flies (in a *w*^*1118*^ background) to identity light-induced changes in lipid levels that were PLC-dependent. Half the control and *norpA*^*P24*^ flies were then exposed to blue light for 5 minutes since the major rhodopsin in the compound eyes (rhodopsin 1) is maximally activated by 480 nm light (Britt *et al*, 1993). The flies were then immediately immersed in liquid nitrogen. We mechanically separated the heads from the bodies by vortexing, and collected them on sieves. We then used whole heads for the following lipidomic analysis since the retina represents a significant proportion (∼20-25%) of the mass of the heads. In addition, ∼90% of PLC activity that is in the heads is due to PLC activity in the retina (Inoue *et al*, 1985). Thus, the vast majority of NORPA-dependent changes in lipid levels is from the retina.

**Figure 1.**
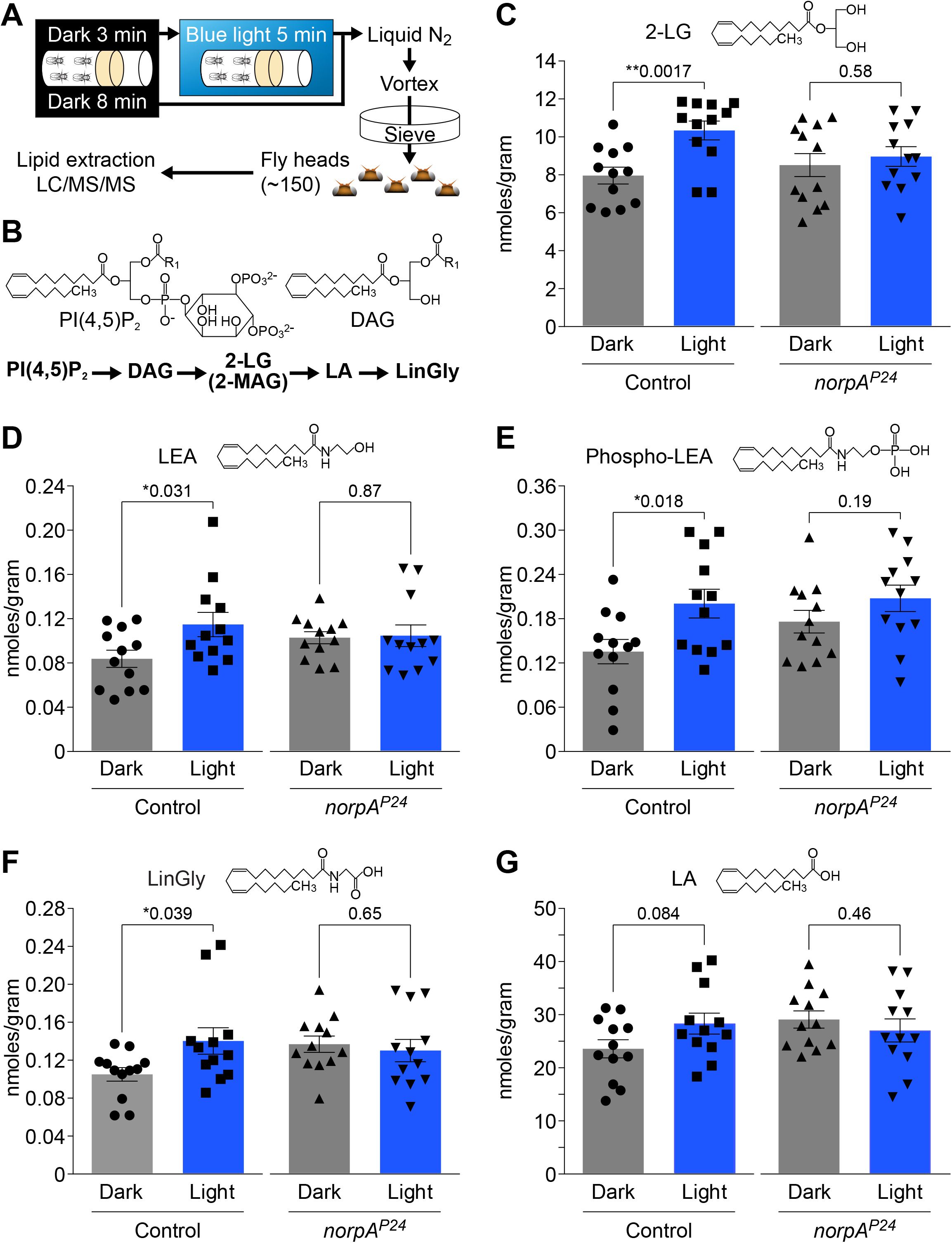
Relative lipid levels in control (*w*^*1118*^) and *norpA*^*P24*^ (in a *w*^*1118*^ background) heads from flies maintained in the dark and after light exposure. **A** Schematic of protocol for collecting heads from flies maintained at 37°C in the dark for 8 minutes or from flies kept in the dark for 3 minutes and then exposed to blue light for 5 minutes. “Dark” is shorthand for flies that were processed using a dim photographic safety light right before the 37°C incubation, which is functionally dark to *Drosophila*. After freezing in liquid N_2_, and vortexing, the heads were collected over a sieve, lipids were extracted and analyzed by LC/MS/MS. **B** Pathway for production of endocannabinoids and other lipids from phosphatidylinositol 4,5-bisphosphate [PI(4,5)P_2_]. DAG, diacylglycerol; 2-LG, 2-linoleoyl glycerol; 2-MAG, 2-monoacylglycerol; LA, linoleic acid. The structures of PI(4,5)P_2_ and DAG are shown. **C—G** Concentrations (nmoles/gram) of the indicated lipids extracted from control and *norpA*^*P24*^ heads that were kept in the dark or exposed to light. All of the lipid metabolites were analyzed from the same set of samples. **(B)** 2-linoleoyl glycerol (2-LG). **(C)** Linoleoyl ethanolamide (LEA). **(D)** Phospho-linoleoyl ethanolamide (phospho-LEA). **(E)** Linoleoyl glycine (LinGly). **(F)** Linoleic acid (LA, C18:2). **Data information:** In (**C—G**), data are presented as mean ± SEM (n=12). *p < 0.05, **p < 0.01 (unpaired Student’s *t*-test).

To quantify the amounts of lipid metabolites in each sample, we used liquid chromatography-tandem mass spectrometry (LC/MS/MS). We analyzed 14 lipids that are produced in *Drosophila* larvae (Tortoriello *et al*, 2013), and which we could reliably identify in *Drosophila* heads in our preliminary studies. These included lipids that are known or could potentially depend on PLC for their biosynthesis since PLC activity is required for the light response. PLC hydrolyzes PIP_2_ to generate IP_3_, H^+^ and DAG. DAG can be metabolized to 2-linoleoyl glycerol (2-LG; Fig 1B) and other 2-monoacylglycerols (2-MAGs). In mammals 2-MAGs such as 2-arachidonoyl glycerol function as endocannabinoids (Gregus & Buczynski, 2020) and recently, 2-LG has been shown to activate and bind to mammalian CB1 when it was ectopically expressed in *Drosophila* (Tortoriello *et al*, 2020). We did not include long PUFAs (C20 and C22) such as arachidonic acid (C20:4) do not appear to be synthesized in *Drosophila* (Shen *et al*, 2010; Tortoriello *et al*., 2013; Yoshioka *et al*, 1985).

Four of the lipids that we characterized displayed significant changes in control fly heads. Most prominent among these four is the endocannabinoid 2-LG (Fig 1C; nmoles/gram: dark 8.0 ± 0.4, light 10.4 ± 0.5). The 30% light-dependent rise in 2-LG levels is highly significant (p=0.0017). Moreover, the light-induced rise in 2-LG that we observed in controls heads did not occur in *norpA*^*P24*^ heads (Fig 1C) demonstrating that the change in 2-LG levels was PLC-dependent. In contrast, to 2-LG, we did not detect significant light dependent changes in two other 2-MAGs analyzed: 2-palmitoyl glycerol (2-PG) and 2-oleoyl glycerol (2-OG; Fig EV1A and B).

Control flies also exhibited a light-dependent increase in the anandamide-related lipid, linoleoyl ethanolamide (LEA; Fig 1D). However, the absolute levels of LEA were ∼100 fold lower than 2-LG (Fig 1C and D). Light did not impact the concentration of LEA in *norpA*^*P24*^ flies (Fig 1D). We also quantified a possible precursor of LEA, phospho-LEA (Liu *et al*, 2006), and found a similar light-dependent rise in phospho-LEA (Fig 1E), which was present at low levels comparable to LEA (Fig 1D). There was a small light-induced increase in phospho-LEA in *norpA*^*P24*^; however, this change was not significant (Fig 1E). In contrast to LEA and phospho-LEA, light did not significantly affect the biosynthesis of other types of *N*-acyl ethanolamides, including those that contained a saturated fatty acid, such as stearoyl ethanolamide (S-EA, C18:0) and palmitoyl ethanolamide (P-EA, C16:0; Fig EV1C and D). We also did not detect a significant light-dependent change in the concentration of oleoyl ethanolamide (O-EA), which is conjugated to the monosaturated fatty acid, oleic acid (OA, C18:1; Fig EV1E). Similarly, light had no impact on OA levels either in control or *norpA*^*P24*^ eyes (Fig EV1F).

The fourth lipid that displayed a light-induced increase in control flies was the NAG— linoleoyl glycine (LinGly; Fig 1B and F). It has been proposed that NAG is produced by conjugation of glycine and fatty acid through fatty acid amide hydrolase (FAAH) (Bradshaw *et al*, 2009). The level of LinGly were unaffected by light in *norpA*^*P24*^ (Fig 1F) indicating that the increase in control flies was PLC dependent (Fig 1B). As with LEA and phospho-LEA, the absolute levels of LinGly were much lower than 2-LG (Fig 1C and F). In contrast to LinGly, the concentrations of all three other *N*-acyl glycine molecules analyzed were not impacted by light (Fig EV1G—I). Thus, all four lipids that displayed light-dependent increases were conjugated to LA. However, linoleic acid (LA) showed only a modest increase in light-stimulated control flies, which was above the threshold for statistical significance (Fig 1G; p=0.084). A previous lipidomics study focusing on PUFAs found that none of the PUFAs analyzed, including LA, changed in the presence of light (Delgado *et al*., 2014). Thus, even though several reports implicate PUFAs as activators of TRP and TRPL (Chyb *et al*., 1999; Delgado & Bacigalupo, 2009; Lev *et al*., 2012), the effects may not be physiologically relevant. Also consistent with previous studies (Shen *et al*., 2010; Yoshioka *et al*., 1985), we did not detect arachidonic acid in any our samples. Taken together, our data indicate that light stimulation promotes biosynthesis of LA-containing endocannabinoids and a NAG, and these increases are all PLC dependent.

### Endocannabinoids and NAG activate TRPL channel *in vitro*

To test whether the linoleoyl conjugates that rise in concentration in response to light increase channel activity *in vitro* we focused on TRPL since TRP is largely retained in the endoplasmic reticulum in tissue cultures and has been refractory to functional analyses. To conduct the current analysis, we used a *Drosophila* cell line (Schneider 2 cells; S2 cells) that contains an integrated *trpl::GFP* gene that can be induced with CuSO_4_ (Parnas *et al*, 2007). Cells that are not exposed to CuSO_4_ do not express *trpl::GFP* and provide a negative control. We introduced a cell permeant, ratiometric Ca^2+^ indicator (Fura-2 AM) into cells and stimulated with different lipids. We then determined the increase in intracellular Ca^2+^ (Ca^2+^_i_) by measuring the change in fluorescence. Finally, we exposed the cells to an ionophore (ionomycin) to determine the maximum possible increase in Ca^2+^, which we calculated by normalizing the maximum value with each treatment relative to the ionomycin response (see Methods).

We focused this analysis on the endocannabinoids (2-LG and LEA) and LinGly, and performed dose response analysis over a 1000-fold concentration range (100 nM—100 μM). We did not include phospho-LEA in these experiments since it is amphipathic and will not flip to the inner leaflet of the plasma membrane when added to the bath solution. Cells that did not express TRPL (Cu^2+^ minus) were unresponsive to 2-LG even at 100 μM 2-LG (Fig 2A and EV2A). In contrast, 2-LG robustly stimulated an increase in Ca^2+^_i_ in cells expressing TRPL (exposed to Cu^2+^) with an EC_50_=5.23 μM (Fig 2A—D). 100 μM 2-LG induced Ca^2+^_i_ that was 51.5 ± 3.3% of the maximum possible value (Fig 2A—C). LEA and LinGly also stimulated an increase in Ca^2+^_i_ in TRPL-expressing cells (EC_50_ μM:

**Figure 2.**
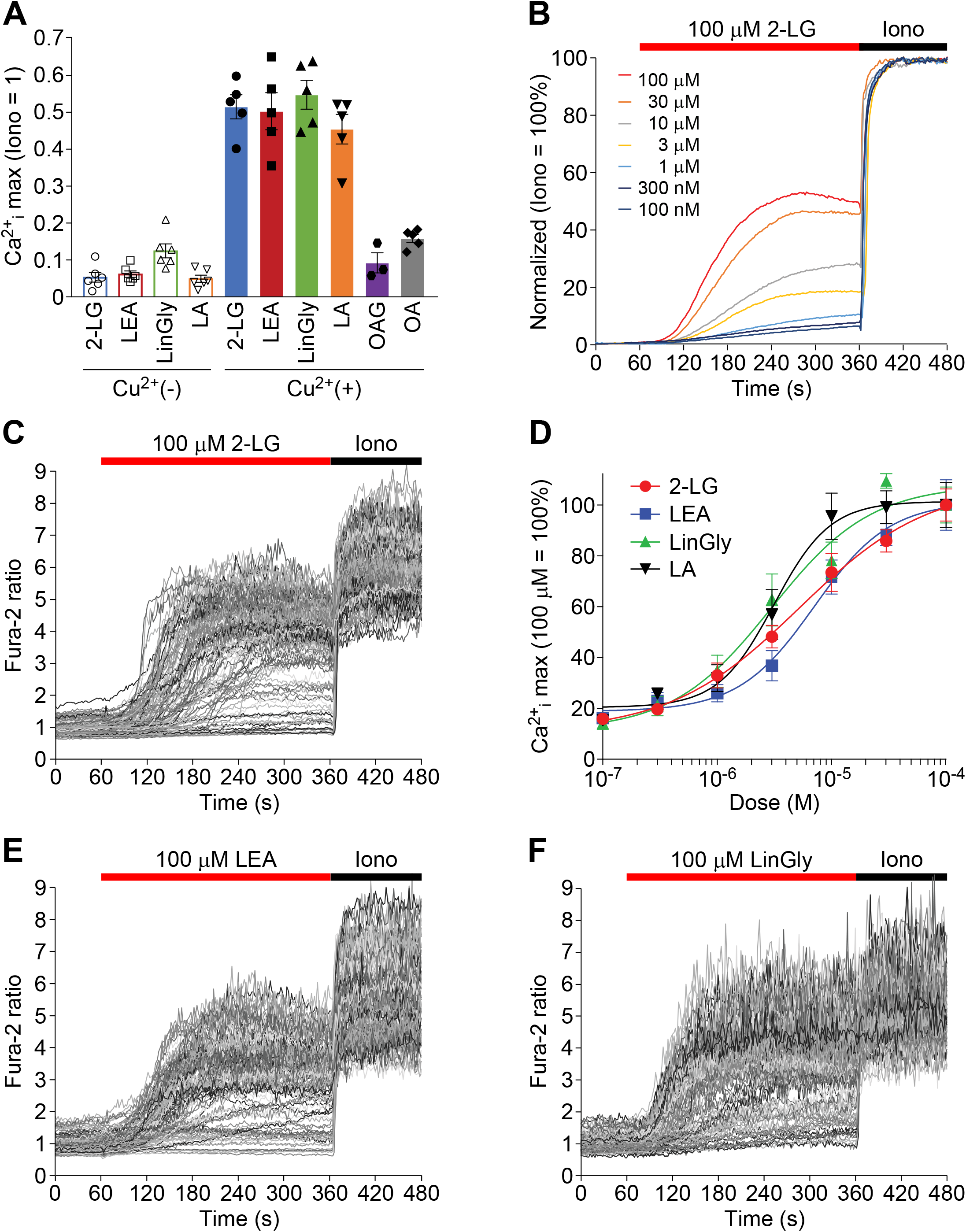
Effects of endocannabinoids and *N*-acyl glycine on TRPL-dependent changes in Fura-2 ratio in S2 cells. **A** Comparison of the maximum increase in intracellular Ca^2+^ (Ca^2+^_i_) in response to the indicated lipids. Lipids were added at 100 μM, except for 300 μM oleic acid (OA). The Cu^2+^ (-) cells did not express TRPL and the Cu^2+^ (+) cells expressed TRPL. **B—F** Fura-2 responses of TRPL-expressing cells to the indicated lipids. The red and black bars **(B, C, E** and **F)** indicate the perfusion of the lipids and the ionomycin (Iono), respectively. **(B)** Dose responses to 2-LG. Values were normalized to Iono. **(C)** Representative traces to 100 μM 2-LG. **(D)** Dose-dependent responses to 2-LG, LEA, LinGly and LA. The data are the maximum Ca^2+^_i_ during the stimulation period (60—360 seconds after addition of the lipids). The basal values were subtracted and the percentages were normalized to the maximum values obtained with 5 μM Iono. The curves were fitted using nonlinear regression with variable slopes. **(E)** Representative traces in response to 100 μM LEA. **(F)** Representative traces in response to 100 μM LinGly. **Data information:** In (**A**), data are presented as mean ± SEM (n=3—6 experiments; ∼100 cells/experiment). In (**D**), data are presented as mean ± SEM (n=5—7; ∼100 cells/experiment).

LEA=7.04, LinGly=2.99; Fig 2A and D—F), but not in Cu^2+^ minus cells, which did not express TRPL (Fig 2A and EV2B and C). In contrast to the efficacy of these linoleoyl conjugates in stimulating a rise in Ca^2+^_i_ in TRPL-positive cells, a membrane-permeable analogue of DAG, 1-oleoyl-2-acetyl-*sn*-glycerol (OAG), was not effective at inducing Ca^2+^_i_ increase even at the highest concentration we tested (Fig 2A and EV2D). OA was also ineffective at activating TRPL (Fig 2A and EV2E) consistent with the lack of increase of OA level upon light stimulation (Fig EV1F).

Since 2-LG, LEA and LinGly contain LA in their structures, channel activation might result from generation of LA either by hydrolysis or degradation. This is plausible since LA activates a TRPL-dependent elevation in Ca^2+^_i_ (Fig 2A and EV2F and G). To assess whether generation of LA from 2-LG, LEA and LinGly activated TRPL, we tested the effects of addition of a MAG lipase inhibitor (JZL 184) and MAG lipase/fatty acid amide hydrolase (FAAH) inhibitor (IDFP) (Long *et al*, 2009; Nomura *et al*, 2008). We found that neither inhibitor reduced Ca^2+^_i_ (Fig 3A), supporting the idea that the endocannabinoids (2-LG and LEA) and NAG promote TRPL activation.

**Figure 3.**
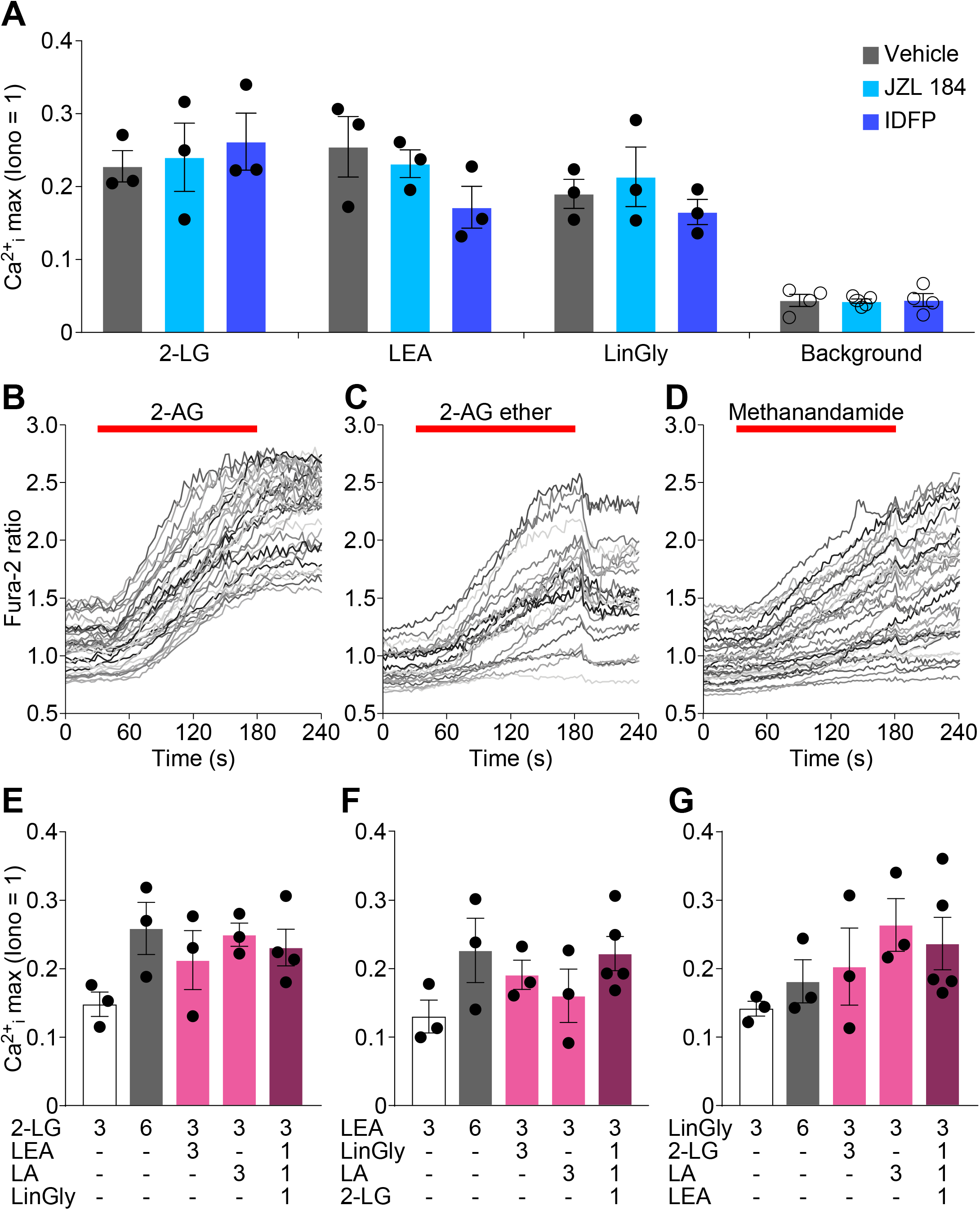
Effects of lipase inhibitors and activation profile of endocannabinoid analogs on Fura-2 responses in TRPL-expressing S2 cells. **A** Testing effects of a monoacyl glycerol lipase (MAGL) inhibitor (JZL 184, 80 nM) and a MAGL/fatty acid amide hydrolase inhibitor (IDFP, 30 nM) on Ca^2+^_i_ by 2-LG (10 μM), LEA (10 μM) and LinGly (10 μM). The cells were pretreated with the inhibitors or the vehicle (0.1% dimethyl sulfoxide) one minute prior to the lipid application. The background Ca^2+^_i_ were obtained in non-induced Cu^2+^ (-) cells with the vehicle alone, 80 nM JZL 184 or 30 nM IDFP. **B—D** Responses of Fura-2 to 100 μM of the indicated lipids. The red bars indicate the stimulation period of the lipids. **E—G** Testing for synergistic or additive effects by mixing combinations of 2-LG, LEA, LinGly and linoleic acid (LA) to TRPL-expressing cells. The numbers indicate the concentration of lipids (μM). **Data information:** In (**A**), data are presented as mean ± SEM (n=3 for TRPL-expressing cells, n=4—5 for background. ∼100 cells/experiment). In (**B—D**), 3 biological repeats were performed (40—50 cells/experiment) for each stimulation. In (**E—G**), data are presented as mean ± SEM (n=3—4; ∼100 cells/experiment).

LA is generated in *Drosophila*, but arachidonic acid is not detectable *in vivo* unless it is supplied in the diet. Nevertheless, we tested the effects of 2-arachidonoyl glycerol (2-AG) and found that it evoked Ca^2+^_i_ increases in TRPL-expressing cells (Fig 3B and EV3A). We tested a stable 2-AG analog, 2-AG ether (Laine *et al*, 2002), which also evoked Ca^2+^_i_ increase (Fig 3C and EV3B). Anandamide (AEA) is similar in structure to LEA. Methanandamide, which is a stable analog of AEA (Abadji *et al*, 1994), also stimulated an increase in Ca^2+^_i_ in TRPL-expressing cells (Fig 3D and EV3C). Thus, arachidonic acid conjugates stimulate TRPL. Since the stable 2-AG analogs activate

TRPL to a comparable extent as 2-AG, this supports the idea that endocannabinoids themselves rather than PUFA metabolites are sufficient to activate TRPL.

2-LG, LEA and LinGly all increase upon light stimulation *in vivo*. Therefore, we tested whether there were synergistic or additive effects resulting from applying mixtures of these lipids. We tested all combinations of 2-LG, LEA, LinGly and LA while maintaining the same total concentration (6 μM). We did not observe either synergistic or additive effects on the increase in Ca^2+^_i_ relative to the same concentration of the single lipids (Fig 3E—G).

### Endocannabinoid acts on TRP and TRPL channels in ommatidia

To address whether the endocannabinoid 2-LG activates TRP and TRPL in photoreceptor cells we isolated ommatidia from flies (Fig EV4A) expressing a genetically encoded Ca^2+^ sensor, GCaMP6f, which is expressed in six out of the eight photoreceptor cells under control of the *rhodopsin 1* (*ninaE*) promotor (*ninaE*>*GCaMP6f*) (Asteriti *et al*, 2017). We performed all analyses in a *norpA*^*P24*^ genetic background to prevent light activation of the TRP and TRPL channels. We stimulated the ommatidia with 2-LG, following by ionomycin to confirm that the ommatidia were viable. An increase in Ca^2+^ was assessed by monitoring the change in fluorescence and dividing it by the basal level of fluorescence (ΔF/F_0_).

We focused this analysis primarily on 2-LG since it is the most abundant lipid that is induced by light. When we applied 2-LG to the bath solution, we observed an increase in Ca^2+^ in *norpA*^*P24*^ photoreceptor cells (Fig 4A, C, G and H, and EV4B). Since the *norpA*^*P24*^ mutation removes the PLC required for phototransduction, the change in fluorescence was not due to light stimulation. We introduced the *norpA*^*P24*^ mutation into a genetic background that removes both TRP and TRPL (*norpA*^*P24*^;*trpl*^*302*^;*trp*^*343*^) and found that most ommatidia showed significantly lower responses to 2-LG (Fig 4B, D, G and H, and EV4C and D). The significance of this reduction (p=0.024) was not due to the two outliers in the *norpA*^*P24*^;+;+ control since the p value would be 7.6 × 10^−4^ in the absence of these two values and the one outlier in the *trp*^*343*^ mutant and two in the *trpl*^*302*^ mutant due to the narrower data distribution. In further support of the conclusion that the TRPC channels are activated by 2-LG *in vivo*, the percentage of no or low responding ommatidia (max ΔF/F_0_ ≤0.2) was significantly higher in the mutant lacking TRPL and TRP (*norpA*^*P24*^;*trpl*^*302*^;*trp*^*343*^; 30.2 ±3.7%) compared with the control (Fig 4I; *norpA*^*P24*^;+;+; 7.8 ±2.0%). Nevertheless, the remaining influx in *norpA*^*P24*^;*trpl*^*302*^;*trp*^*343*^ flies could be due to the Na^+^/Ca^2+^ exchanger (CalX), which we can run in reverse in fly photoreceptor cells (Wang *et al*, 2005b) or potentially to lipid modulation of voltage-gated channels (Elinder & Liin, 2017).

**Figure 4.**
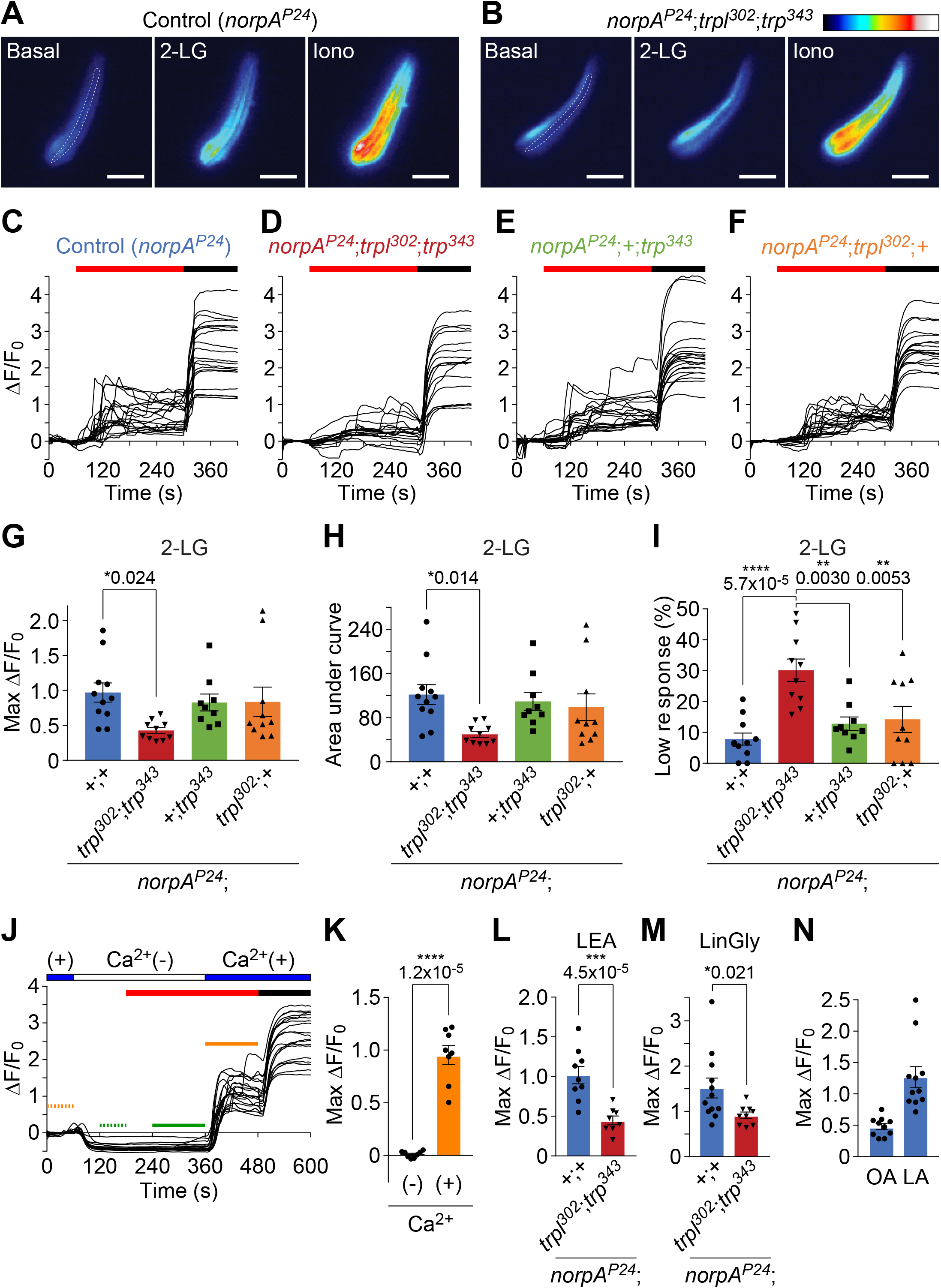
Monitoring responses of photoreceptor cells stimulated with endocannabinoids and N-acyl glycine with GCaMP6f. **A, B** Representative GCaMP6f responses to 2-LG in control ommatidia (*norpA*^*P24*^) and in *norpA*^*P24*^;*trpl*^*302*^;*trp*^*343*^ ommatidia. The ommatidia were stimulated with 30 μM 2-LG, followed by 5 μM ionomycin (Iono) to confirm the responsiveness of the GCaMP6f in the photoreceptor cells. The changes in fluorescence are shown in pseudo colors (0-255). The dotted lines in the images of the basal conditions (before addition of 2-LG) outline individual rhabdomeres. The 2-LG images were obtained at the 300 seconds time points in **C and D**. Scale bars, 20 μm. **C—F** Traces showing representative Ca^2+^_i_ responses (ΔF/F_0_) in photoreceptor cells from the indicated flies. The red and black bars indicate application of 30 μM 2-LG (60— 300 seconds) and 5 μM ionomycin (after 300 seconds), respectively. **(C)** Control (*norpA*^*P24*^). **(D)** *norpA*^*P24*^;*trpl*^*302*^;*trp*^*343*^. **(E)** *norpA*^*P24*^;+;*trp*^*343*^. **(F)** *norpA*^*P24*^;*trpl*^*302*^;+. **G** ΔF/F_0_ indicates the maximum Ca^2+^_i_ responses during the stimulation period (4 minutes: 60—300 seconds in **C—F**) divided by the basal fluorescence levels. **H** Quantification of area under curve during the stimulation period (4 minutes: 60—300 seconds in **C—F**). **I** Proportion of no or low responding photoreceptor cells (max ΔF/F_0_ ≤0.2) during the stimulation period (4 minutes: 60—300 seconds in **C—F**). **J** 2-LG stimulated Ca^2+^ influx rather than Ca^2+^ release from internal stores in isolated ommatidia. The blue bars near the top indicate the bath that contained 1.5 mM Ca^2+^. The white bar indicates the bath with no addition of Ca^2+^. The red and black bars indicate application of 30 μM 2-LG and 5 μM ionomycin, respectively. The dotted and solid orange lines indicate ΔF/F_0_ in a Ca^2+^-containing bath in the absence (dotted) and presence (solid) of 2-LG. The dotted and solid green lines indicate ΔF/F_0_ in a bath without added Ca^2+^ in the absence (dotted) and presence (solid) of 2-LG. **K** Quantification of the maximum Ca^2+^_i_ responses to 30 μM 2-LG in the absence (-) or the presence (+) of 1.5 mM extracellular Ca^2+^. ΔF/F_0_ in each Ca^2+^ condition was calculated using values in the periods indicated by green and orange dotted and solid lines in **(J)**. **L, M** GCaMP6f responses of control (*norpA*^*P24*^) and *norpA*^*P24*^;*trpl*^*302*^;*trp*^*343*^ photoreceptor cells to 100 μM LEA or 30 μM LinGly as indicated. The maximum Ca^2+^_i_ responses during the stimulation period (4 minutes) were used for the calculations. **N** GCaMP6f responses of control (*norpA*^*P24*^) photoreceptor cells to 30 μM oleic acid (OA) and 30 μM linoleic acid (LA) as indicated. The maximum Ca^2+^_i_ responses during the stimulation period (4 minutes) were used for the calculations. **Data information:** In (**G, H**), data are presented as mean ± SEM. n=9—11 experiments; 13—30 ommatidia/experiment. *p < 0.05. One-way ANOVA with Dunnett’s *post hoc* analysis. In (**I**), the data are presented as mean ± SEM. n=9—11 experiments; 13—30 ommatidia/experiment. **p < 0.01, ****p < 0.0001. One-way ANOVA with Tukey’s *post hoc* analysis. In (**K**), the data are presented as mean ± SEM. n=8. 13—30 ommatidia/experiment. ****p < 0.0001. Paired Student’s *t*-test. In (**L, M**), the data are presented as mean ± SEM. n=9—11. 13—30 ommatidia/experiment. *p < 0.05, ***p < 0.001. Unpaired Student’s *t*-test. In (**N**), the data are presented as mean ± SEM. n=10— 11. 14—30 ommatidia/experiment.

The reduction in fluorescence (ΔF/F_0_) in the *norpA*^*P24*^;*trpl*^*302*^;*trp*^*343*.^ mutant demonstrates that the rise in ΔF/F_0_ was due primarily to TRP and TRPL. Elimination of just TRP or TRPL resulted in only small differences from the *norpA*^*P24*^ control, which were not statistically significant (Fig 4E—I). The minimal changes upon elimination of just TRP or TRPL is consistent with analyses using electrophysiological studies showing that loss of TRPL alone has virtually no effect on the amplitude of the light response (Niemeyer *et al*., 1996), while removal of TRP has only minimal effects on the response amplitude under dim or moderate light conditions (Minke, 1982; Minke *et al*, 1975). The 2-LG induced an increase in ΔF/F_0_ only in the presence and not the absence of Ca^2+^ in the bath solution (Fig 4J and K) demonstrating that the increase in GCaMP6f fluorescence was due to Ca^2+^ influx and not to release of Ca^2+^ from internal stores.

We also examined whether LEA and LinGly stimulated a rise in Ca^2+^_i_. We found that these lipids induced Ca^2+^_i_ increases in the *norpA*^*P24*^ control, which were significantly higher than those in the *norpA*^*P24*^;*trpl*^*302*^;*trp*^*343*^ mutant (Fig 4L and M). In the absence of the one outlier for LinGly, the significance is essentially unchanged (p=0.020).

Consistent with previous data indicating that LA but not OA is effective in activating the TRPC channels (Chyb *et al*., 1999; Delgado & Bacigalupo, 2009), we found that LA activated ommatidia at a level similar to that of 2-LG, whereas OA only evoked the minimum Ca^2+^ increase similar to that observed in ommatidia lacking TRPL and TRP (Fig 4N).

## Discussion

Activation of PLC following light stimulation is critical for opening of the TRP and TRPL channels in photoreceptor cells. However, many lipids could potentially be generated following stimulation of PLC. To identify candidate lipids that modulate these TRPC channels *in vivo*, we set out to identify lipids that increase in response to light.

Since phototransduction is dependent on the PLC encoded by *norpA*, we designed the analysis to find lipids that increased upon light stimulation in wild-type but in not in the *norpA* mutant. We found that wild-type but not *norpA* mutant flies exhibit light-dependent increases in the endocannabinoids 2-LG, LEA, phospho-LEA as well as the NAG (LinGly), and that 2-LG, LEA and LinGly activate TRPC channels *in vitro* and in isolated ommatidia. The results suggest that one or more of these lipids activates the TRP and TRPL channels in photoreceptor cells.

We suggest that the endocannabinoid 2-LG is the relevant lipid that activates the TRPC channels *in vivo*. 2-LG is generated at levels that are ∼60—100-fold higher than LEA, phospho-LEA or LinGly. In addition, the light-dependent rise in 2-LG was more significant (p=0.0017) than the other three lipids (p=0.018—0.039). Moreover, the statistical significance of LEA and LinGly depended on the one and two outliers, respectively. In contrast to the highly significant light-dependent rise in 2-LG in control flies, the levels of 2-LG in either dark or light exposed *norpA*^*P24*^ heads were virtually identical to the control maintained in the dark. We suggest that enough 2-LG is generated to activate the TRPC channels since the precursor for 2-LG (DAG) is estimated to be produced at near millimolar levels (Raghu & Hardie, 2009), and 30 μM 2-LA is sufficient to activate the TRPC channels in isolated ommatidia. The TRPC channels in the rhabdomeres are activated within 20 milliseconds (Ranganathan *et al*, 1991), and our lipidomics analysis was performed following 5 minutes of illumination. Although technically challenging, in the future it would be of interest to assess whether the rise 2-LG can be detected over a much shorter time frame.

In further support of the model that 2-LG is the physiologically relevant activator of the TRPC channels, mutation of *Drosophila inaE*, which encodes a DAG lipase necessary for the sn-1 hydrolysis for production of 2-MAG from DAG, causes a transient ERG phenotype that resembles the *trp* mutant (Leung *et al*, 2008). The *inaE* mutant phenotype indicates that one or more lipid metabolites produced subsequent to light-dependent production of DAG contributes to TRP channel activation. However, this work did not clarify whether the relevant lipid is 2-LG or some other metabolite such LA. We suggest that the *inaE* phenotype is not as severe as the *norpA*^*P24*^ null mutant since the viable *inaE* alleles are hypomorphic (Leung *et al*., 2008). Several studies found that LA and other PUFAs can activate the TRPC channels (Chyb *et al*., 1999; Delgado & Bacigalupo, 2009; Lev *et al*., 2012). We also find that LA can activate the TRPC channels. However, a previous report concludes that PUFAs including LA remain unchanged upon illumination (Delgado *et al*., 2014). This is consistent with our lipidomic analysis indicating that the modest rise in LA in light stimulated animals is not significant.

While LEA, phospho-LEA and LinGly are also increased in a light-dependent manner and are effective at activating the TRPC channels, the findings that the light-dependent increases are much lower than 2-LG suggests that they are not likely to be the prime lipids that activate the highly abundant TRP and TRPL channels in photoreceptor cells. We suggest that the endocannabinoids (LEA and phospho-LEA) and the NAG (LinGly) do not function primarily in activation of TRP and TRPL, but rather in some other light-dependent function in photoreceptor cells. One possibility is that these lipids might regulate synaptic vesicle recycling in photoreceptor cells, since this dynamic process depends on proper lipid content at the synapse (Marza *et al*, 2008). Moreover, if flies are fed a diet deficient in PUFA, this causes a deficit in the on- and off-transient responses in the electroretinogram, which reflects a decrease in signal transmission from photoreceptor cells to their postsynaptic partners (Ziegler *et al*, 2015). Although increases of LEA, phosphor-LEA and LinGly are PLC-dependent, it is unclear how these lipids are produced in *Drosophila*.

Our results indicate that the specific fatty acid conjugate and the degree of saturation of the fatty acid in the lipid are factors that contribute to activation of TRP and TRPL. All of the lipids that increased in a light- and NORPA-dependent manner were conjugated to the PUFA, linoleic acid (C18:2). In contrast the levels of two other 2-MAGs analyzed that included either a saturated fatty (palmitic acid, C16:0) or a monounsaturated fatty acid (oleic acid, C18:1) were not increased by light. Moreover, oleic acid was ineffective at activated TRPL *in vitro* or in increasing Ca^2+^ responses in ommatidia.

An open question concerns the mechanism through which 2-LG activates the channels. We propose that the endocannabinoid, 2-LG, directly activates TRP and TRPL *in vivo*. In mammals, plant-derived cannabinoids and endocannabinoids bind to the G-protein coupled receptors CB1 and CB2 (Gregus & Buczynski, 2020). However, there are no *Drosophila* CB1 or CB2 homologs (McPartland *et al*., 2001). At least six mammalian TRP channels are activated *in vitro* by cannabinoids and endocannabinoids, including four TRPV channels, TRPA1 and TRPM8 (Bradshaw *et al*, 2013; Muller *et al*, 2019). An interaction between rat TRPV2 and cannabidiol was recently identified in a cryo-EM structure (Pumroy *et al*, 2019) and differences in the putative binding sites among mammalian TRP channels has been modeled and discussed (Muller *et al*, 2020; Muller & Reggio, 2020). Similarly, the *Drosophila* TRPC channels, TRP and TRPL, may also be receptors for endocannabinoids. Since CB1 and CB2 are not present in *Drosophila* (McPartland *et al*., 2001), we suggest that TRP channels comprise a class of ionotropic cannabinoid receptors conserved from flies to humans.

A previous and highly intriguing study proposed that the TRP and TRPL channels are mechanically-gated following light-induced activation of the phototransduction cascade and stimulation of NORPA (Hardie & Franze, 2012). Their concept is that following stimulation of PLC, the DAG that remains in the membrane is smaller than PIP_2_, thereby resulting in a conformational change in the plasma membrane that causes mechanical activation of the channels. We suggest the model that allosteric modulation of TRP and TRPL by 2-LG along with conformational changes in the membrane due to hydrolysis of PIP_2_ both contribute to activation of the TRPC channels. This dual mechanism would ensure that production of 2-LG alone, or conformation changes of the membrane alone would be insufficient to activate TRP and TRPL. Since generation of 2-LG or movements in the plasma membrane could in principle occur without stimulation of PLC, this dual mechanism would reduce the probability of channel activation independent of light. It has also been posited that mechanical stimulation in combination with protons produced from PIP_2_ hydrolysis could collaborate to promote TRPC channel activation (Hardie & Franze, 2012). A requirement for two PLC-dependent changes to activate TRP and TRPL would have the great benefit of minimizing noise in photoreceptor cells, which is essential for photoreceptor cells to achieve their exquisite single photon sensitivity.

## Materials and methods

### Sources of fly stocks and rearing

*w*^*1118*^ was used as the wild-type control. The following flies were obtained from the Bloomington Stock Center (stock numbers are indicated): *trp*^*343*^ (#9046), *trpl*^*302*^ (#31433).

The following stocks were provided by the indicated investigators: *cn*^*1*^,*trpl*^*302*^,*bw*^*1*^;*trp*^*343*^,*ninaE-GCaMP6f*/*Tb*^*1*^ (R. Hardie) and *norpA*^*P24*^ (W. Pak), which was outcrossed to *w*^*1118*^. The *w*^*1118*^,*norpA*^*P24*^ flies were used throughout this work but are referred to as *norpA*^*P24*^ for brevity. Flies were reared on standard cornmeal-yeast media: 24,900 ml distilled water, 324 g agar (66-103, Genesee scientific), 1,800 g cornmeal (NC0535320, lab scientific), 449 g yeast (ICN90331280, MP Biomedicals), 240 ml Tegosept (30% in ethanol; H5501, Sigma Aldrich), 72 ml propionic acid (81910, Sigma Aldrich), 8.5 ml phosphoric acid (438081, Sigma Aldrich) and 2,400 ml molasses (62-118, Genesee Scientific). Flies were initially raised in vials or bottles containing the media at 25°C in a chamber under 12-hour light/12-hour dark cycle and transferred to 24-hour dark conditions before experiments as indicated below.

### Chemicals

The following chemicals were obtained from Cayman Chemical: linoleic acid (#90150), 2-linoleoyl glycerol (#62260), linoleoyl ethanolamide (#90155), linoleoyl glycine (#9000326), oleic acid (#9000326), 1-oleoyl-2-acetyl-*sn*-glycerol (OAG, #62600), 2-arachidonoyl glycerol (#62160), 2-arachidonoyl glycerol ether (#62165), R-1 methanandamide (#90070), JZL 184 (#13158) and IDFP (#10215). The chemicals were dissolved in ethanol or DMSO and kept at −80°C.

### Exposing flies to light and collecting fly heads for lipidomic analyses

Bottles containing flies were transferred to and maintained in the dark for 7 days after egg laying and handled under a dim red photographic safety light throughout the experiments. ∼150 flies (0—4 days old) were collected and transferred into bottles containing fly food. The bottles were wrapped with aluminum foil and placed in the dark. After two days, flies were starved in the dark by transferring the flies into bottles containing 1% agarose, wrapped with aluminum foil and placed in the dark. After 15— 17 hours, the flies were anesthetized with CO_2_ and transferred into 50 mL tubes (352070, BD Falcon). After 1—2 minutes when the flies began to move, we plugged each tube with a cotton ball, which we pushed down to the 10-mL line. The tubes were covered with aluminum foil and placed in a rack for 10 minutes to allow the flies to continue to recover from the CO_2_ exposure. After removing the aluminum foil, we left the cotton ball in each tube and secured the top of each tube with a screw cap. We then placed the tubes in a 37°C incubator for 3 minutes since PLC activity is higher at 37°C than at the standard incubation temperature of 25°C (Huang *et al*., 2004).

To enable us to compare lipids that increase upon light exposure, we either maintained the tubes with the flies in the dark or exposed the flies to blue light from the side of the tube (∼0.3—1.0 mW at the distal and the proximal sides of the tube, respectively) for 5 minutes. The tubes were immediately immersed in liquid nitrogen for 30 seconds. To mechanically separate the head and bodies, we removed the cotton plugs, reinserted the screw caps, and vigorously vortexed the tubes. The frozen samples were then passed through 25 and 40 mesh sieves. The fly heads, which were trapped on the 40 mesh sieves, were quickly transferred into chilled 1.5 ml black microfuge tubes (T7100BK, Argos technologies) and stored at −80°C until we performed the lipid extractions.

### Lipid extraction from fly heads

500 μL of methanol was added to each tube containing 4—8 mg of fly heads followed by 100 pmols, deuterium-labeled *N*-arachidonoyl glycine (d8NAGly), to act as an internal standard. The tubes were then closed, vortexed for 30 seconds and left in darkness on ice for 1 hour, vortexed again for 30 seconds and left to process for another hour in darkness on ice. The samples were then centrifuged at 19,000xg at 24°C for 20 minutes. The supernatants were collected and placed in polypropylene tubes (15 ml) and 1.5 ml of HPLC-grade water was added making the final supernatant/water solution 25% organic. A partial purification of lipids was achieved using a preppy apparatus assembled with 500 mg C18 solid-phase extraction columns. The columns were conditioned with 5 mL of HPLC-grade methanol immediately followed by 2.5 mL of HPLC-grade water under pressure. The supernatant/water solution was then loaded onto the C18 column and washed with 2.5 mL of HPLC grade water followed by 1.5 mL of 40% methanol. Elutions of 1.5 mL of 60%, 70%, 85% and 100% methanol were collected in individual autosampler vials and then stored in a −80°C freezer until mass spectrometer analysis.

### LC/MS/MS analysis and quantification

Samples were removed from the −80°C freezer and allowed to warm to room temperature then vortexed for approximately 1 minute before being placed into the autosampler and held at 24°C (Agilent 1100 series autosampler, Palo Alto, CA) for the LC/MS/MS analysis. 20 μL of eluants were injected separately to be rapidly separated using a C18 Zorbax reversed-phase analytical column to scan for individual compounds.

Gradient elution (200 μL/min) then occurred, under the pressure created by two Shimadzu 10AdVP pumps (Columbia, MD). Next, electrospray ionization was done using an Applied Biosystems/MDS SCIEX (Foster City, CA) API3000 triple quadrupole mass spectrometer. All compounds were analyzed using multiple reaction monitoring (MRM). Synthetic standards were used to generate optimized MRM methods and standard curves for analysis. We reported the MRM parent/fragment pairs previously (Tortoriello *et al*, 2013), with the exception of phosphoLEA, which is MRM[-] 403.5/58.5. The mobile phases are also the same as we reported previously (Tortoriello *et al*, 2013): mobile phase A, 80% HPLC-grade H_2_O/20% HPLC-grade methanol, 1 mM ammonium acetate; mobile phase B, 100% HPLC-grade methanol, 1 mM ammonium acetate.

### Ca^2+^ imaging

Schneider 2 (S2) cells carrying a *trpl-egfp* transgene (gift from B. Minke) (Parnas *et al*., 2007) were grown in 60 mm dishes (353002, Falcon) at 25°C in 4 mL Schneider’s media (21720-024, Gibco) that contained 10% inactivated fetal bovine serum (10437-028, Gibco) and 50 units/mL penicillin/streptomycin (15140-122, Gibco). S2 cells were seeded on 8 mm round cover glasses (Matsunami) in a 35 mm dishes (1000-035, Iwaki). We then added 2 μL 500 mM CuSO_4_ (039-04412, Wako) to 2 mL culture medium (final 500 μM) and incubated the cells for 24 hours in a 25°C incubator to induce expression of the gene encoding TRPL::EGFP. The cells without the CuSO_4_ treatment were used as the control cells that did not express TRPL. To load the cells with Fura-2 AM, we added a 1 mL of the following mixture to the culture media and incubated the cells at 25°C incubator for 1—3 hours: 5 μM Fura-2 AM (F-1201, Life Technologies), 250 μM probenecid (162-26112, Wako), 20% pluronic F-127 (P2443, Sigma). The cells were then allowed to recover for 15 minutes in a bath solution containing 130 mM NaCl, 5 mM KCl, 2 mM MgCl_2_, 2 mM CaCl_2_, 30 mM sucrose, and 10 mM N-Tris(hydroxymethyl)methyl-2-aminoethanesulfonic acid (TES), after adjusting the pH to 7.2 with NaOH. The cover glasses were mounted in a chamber (RC-26G; Warner Instruments) connected to a gravity flow system to deliver various stimuli. A xenon lamp was used as an illumination source. To obtain fluorescent intensities of Ca^2+^-bound and Ca^2+^-free Fura-2, we excited the cells at 340 and 380 nm, respectively, and emission was monitored at 510 nm with a sCMOS camera (Zyla 4.2 Plus; Andor Technology) used with a fluorescent microscope (Eclipse TE2000-U, Nikon). The data were acquired using iQ2 software (Andor Technology) and the ratio values (340/380) were calculated with Fiji software (Schindelin *et al*, 2012).

We obtained an average trace for each sample and calculated the change in Ca^2+^_i_ as follows: Ca^2+^_i_ response=(F_res_ – F_min_)/(F_max_ – F_min_). To normalize the responses, we subtracted the minimum values during the basal period (F_min_) from the responses every 3 sec (F_res_). We also subtracted the F_min_ from the maximum value obtained due to addition of the ionomycin (F_max_) (I0634, Sigma). After the normalization we extracted the maximum increase in Ca^2+^_i_ (Ca^2+^_i_ max) during the stimulation period for further analysis. During stimulation with 2-AG ether, we observed non-specific Ca^2+^_i_ responses, which we recognized due to stochastic and sudden ratio increases in CuSO_4_-induced (TRPL-expressing) and non-induced control cells. Traces containing these non-specific responses were omitted from the analyses.

### Dissociation of ommatidia

All ommatidia were isolated from *norpA*^*P24*^ flies to prevent light-induced activation of the TRP and TRPL channels. The flies were maintained under standard light/dark cycles and transferred to a constant darkness after initiating new crosses to eliminate any possibility of light-induced retinal degeneration. Male progeny were selected for the experiments with isolated ommatidia because *norpA* is on the X chromosome, thereby simplifying the crosses needed to obtain flies with the *norpA*^*P24*^ mutation, which is recessive. The *trpl*^*302*^ (Niemeyer *et al*., 1996) and *trpl*^*343*^ (Wang *et al*, 2005a) mutations are recessive. The control flies were heterozygous for *trpl*^*302*^ and *trpl*^*343*^ (*norpA*^*P24*^/*Y*;*cn*^*1*^,*trpl*^*302*^,*bw*^*1*^/*+*;*trp*^*343*^,*ninaE-GCaMP6f*/*+*) and were obtained by crossing *norpA*^*P24*^ females and *cn*^*1*^,*trpl*^*302*^,*bw*^*1*^;*trp*^*343*^,*ninaE-GCaMP6f*/*Tb*^*1*^ males. The *trp*^*343*^ mutant was also heterozygous for *trpl*^*302*^ (*norpA*^*P24*^/*Y*;*cn*^*1*^,*trpl*^*302*^,*bw*^*1*^/*+*;*trp*^*343*^,*ninaE-GCaMP6f*/*trp*^*343*^) and was obtained by crossing *norpA*^*P24*^;+;*trp*^*343*^ females and *cn*^*1*^,*trpl*^*302*^,*bw*^*1*^;*trp*^*343*^,*ninaE-GCaMP6f*/*Tb*^*1*^ males. The *trpl*^*302*^ mutant was also heterozygous for *trp*^*343*^ (*norpA*^*P24*^/*Y*;*cn1,trpl*^*302*^,*bw*^*1*^;*trp*^*343*^,*ninaE-GCaMP6f*/*+*) and was obtained by crossing *norpA*^*P24*^;*trpl*^*302*^ females and *cn*^*1*^,*trpl*^*302*^,*bw*^*1*^;*trp*^*343*^,*ninaE-GCaMP6f*/*Tb*^*1*^ males. The *trpl*^*302*^;*trp*^*343*^ mutant (*norpA*^*P24*^/*Y*;*cn*^*1*^,*trpl*^*302*^,*bw*^*1*^;*trp*^*343*^,*ninaE-GCaMP6f*/*trp*^*343*^) was obtained by crossing *norpA*^*P24*^;*trpl*^*302*^;*trp*^*343*^ females and *cn*^*1*^,*trpl*^*302*^,*bw*^*1*^;*trp*^*343*^,*ninaE-GCaMP6f*/*Tb*^*1*^ males.

Dissection of ommatidia was performed similar to that described previously (Hardie, 1991). Briefly, we performed dissections under a dim LED light source with a red filter (RG610, Schott), which is functionally equivalent to darkness for the flies. To conduct each experiment, two males (within 4 hours of eclosion) were collected using CO_2_ and the heads were removed, briefly soaked in 70% ethanol and then immersed in a drop of dissection media containing Schneider’s medium and 0.2% bovine serum albumin (fatty acid-free, A8806, Sigma). The four eye cups were dissected using forceps and the retina were scooped out using a micro scooper made from a minutien pin (26002-10, Fine Science Tools). The retinas were collected using a fire-polished trituration glass pipette made from a glass capillary (outer diameter 1.2 mm, inner diameter 0.69 mm, GC120-10, Warner Instruments) and washed with and incubated in fresh dissection media for 20 minutes in the dark. Surrounding pigmented glia were removed by rapid aspiration/expiration and the retina were transferred to a drop (30 μL) of the dissection media. Ommatidia were then mechanically dissociated by repetitive pipetting using three fire-polished trituration pipettes with different inner diameters until almost all the ommatidia were isolated from the lamina layers. Dissociated ommatidia were immediately used for subsequent imaging experiments and maintained in the drop in a dark for up to 60 minutes.

### GCaMP6 imaging

Each ommatidial suspension (8—9 μL) was placed on the glass bottom of a chamber (RC-26G; Warner Instruments). Cells were allowed to settle down to the bottom for 3—4 minutes. To wash out floating cells, the chamber was filled and perfused with an extracellular solution containing 120 mM NaCl, 5 mM KCl, 4 mM MgCl_2_, 1.5 mM CaCl_2_, 25 mM L-proline, 5 mM L-alanine, and 10 mM TES, which was adjusted to pH 7.15 with NaOH. The Ca^2+^-free experiments were performed with a solution that was nominally Ca^2+^-free by omitting 1.5 mM CaCl_2_ from the extracellular solution. A xenon lamp was used as an illumination source.

To monitor the fluorescent intensity of the GCaMP6f, the ommatidia were excited with 472 nm light and emissions were monitored at 520 nm with a sCMOS camera (Zyla 4.2 Plus; Andor Technology) attached to a fluorescent microscope (Eclipse TE2000-U, Nikon). To minimize photobleaching of GCaMP6f and an exhaustion of cells caused by light activation of the rhodopsins, we excited the ommatidia every 6 seconds for 60 milliseconds. Ionomycin (5 μM) was applied in the end of protocol to evaluate the viability of ommatidia.

Data were acquired with iQ2 software (Andor Technology) and the fluorescent intensities were calculated with Fiji software (Schindelin *et al*., 2012). A region of interest (ROI) was defined as the distal half (the outer side) of the ommatidia since Ca^2+^_i_ responses were relatively higher in this region (Fig 5A and B) and the proximal half (the inner side) of ommatidia was prone to vibration during the perfusion. Typically, 15—30 ommatidia in a field was chosen for analysis and cells with obvious damage or small responses to ionomycin or were out of focus were omitted from the analysis. Changes in fluorescence intensity (ΔF/F_0_) was used to assess the Ca^2+^_i_ responses [(F_t_ – F_basal_)/F_basal_]. F_t_ corresponds to the value obtained every 6 seconds. F_basal_ is the average during the first 1 minute (0.1% EtOH alone) in every ommatidium. The background values were measured in *norpA*^*P24*^ ommatidia that do not express GCaMP6f in the presence of 0.1% EtOH. The average background values were subtracted from the fluorescent intensities in all samples. The maximum response (max ΔF/F_0_) with each lipid (in 0.1% EtOH) was obtained during the 4-minute stimulation period (60—300 seconds) after addition of the lipids. The areas under the curve during the stimulation period (60—300 seconds after addition of the lipids) were calculated using a trapezoidal rule [(F_t_+F_t+1_)/2 × 6 (sampling interval)]. For no or low responding ommatidia, the number of cells having max ΔF/F_0_ ≤0.2 were counted and divided by the total number of cells in each sample to obtain the proportion.

### Quantification and statistical analysis

The data are represented as means ± SEM. The number of repeated times for each experiment (n) is indicated in the figure legends. We used the unpaired, two-tailed Student’s *t*-test to determine the statistical significance of two samples that had equal variance. In experiments in which we compared two sets of data that did not have equal variance, we used Welch’s *t*-test. To determine the statistical significance of the data using ommatidia in Ca^2+^-free versus Ca^2+^-containing bath conditions, we used the paired, two-tailed Student’s *t*-test. To evaluate the statistical significance of multiple samples, we used one-way ANOVA with Dunnett’s *post hoc* analysis. To evaluate the statistical significance of multiple samples in the no or low response population, we used one-way ANOVA with Tukey’s *post hoc* analysis. Statistical tests were performed using Prism 7 (Graphpad). Asterisks indicate statistical significance, where *p < 0.05, **p < 0.01, ***p < 0.001 and ****p < 0.0001.

## Data Availability

No data that requires deposition in a public database.

## Acknowledgments

This work was supported by a grant to C.M. from the National Eye Institute (EY010852), a grant to H.B. from the National Institute on Drug Addiction (DA039463), a grant to M.T. from a Grant-in-Aid for Scientific Research from the Ministry of Education, Culture, Sports, Science and Technology in Japan (#15H02501), and grants to T.S. from a Grant-in-Aid for Scientific Research from the Ministry of Education, Culture, Sports, Science and Technology in Japan (#17H07337 and #18K06495). We thank Dr. Baruch Minke (Hebrew University) for sharing the *trpl*-expressing stable S2 cells, and Dr. Roger Hardie (Cambridge University) for sharing valuable fly stocks and suggestions for the ommatidia dissociation protocol.

## Author Contributions

Conceptualization, T.S. and C.M.; Methodology, T.S., H.B.B., E.L., and C.M.; Investigation, T.S., H.B.B., and E.L.; Formal Analysis, T.S., H.B.B., and E.L.; Writing — Original Draft, T.S. and C.M.; Writing — Review and Editing, T.S., H.B.B., M.T., and C.M.; Funding Acquisition, T.S., H.B.B., M.T., and C.M.; Supervision, T.S. and C.M.

## Expanded View Figure legends

**Figure EV1**. Relative lipid levels in control (*w*^*1118*^) and *norpA*^*P24*^ (in a *w*^*1118*^ background) heads from flies maintained in the dark and after light exposure.

**A—I** All these lipid metabolites were measured (nmoles/gram) in the same set of samples used in *Figure 1*. **(A)** 2-palmitoyl glycerol (2-PG). **(B)** 2-oleoyl glycerol (2-OG).

**(C)** Stearoyl ethanolamide (S-EA). **(D)** Palmitoyl ethanolamide (P-EA). **(E)** Oleoyl ethanolamide (O-EA). **(F)** Oleic acid (OA). **(G)** Stearoyl glycine (S-Gly). **(H)** Palmitoyl glycine (P-Gly). **(I)** Oleoyl glycine (O-Gly).

**Data information:** Data are presented as mean ± SEM. n=12. Unpaired Student’s *t*-tests.

**Figure EV2**. Effects of linoleoyl-conjugates and linoleic acid on S2 cells in which *trpl* was not induced [Cu^2+^ (-), **(A—C** and **G)**] or *trpl* was induced with Cu^2+^ **(D—F)**.

The cells were loaded with Fura-2 AM and each lipid (100 μM or 300 μM) was applied exogenously by perfusion. Ionomycin (Iono; 5 μM) was used to confirm cell viability. The cells were excited at 340 nm and 380 nm to obtain the Fura-2 ratio. The red and black bars indicate the addition of the lipids or Iono, respectively. **(A)** 2-LG, Cu^2+^ (-). **(B)** LEA, Cu^2+^ (-). **(C)** LinGly, Cu^2+^ (-). **(D)** OAG, Cu^2+^ (+). **(E)** OA, Cu^2+^ (+). **(F)** LA, Cu^2+^ (+). **(G)** LA, Cu^2+^ (-).

**Data information:** 3 or more independent assays were performed for each stimulation (∼100 cells/experiment).

**Figure EV3**. Effects of endocannabinoid analogs on S2 cells in which *trpl* was not induced [Cu^2+^ (-)].

The red bars indicate the addition of 100 μM of the indicated lipids by perfusion. The cells were excited at 340 nm and 380 nm to obtain the Fura-2 ratio. **(A)** 2-arachidonoyl glycerol (2-AG). **(B)** 2-AG ether. **(C)** Methanandamide.

**Data information:** 3 or more independent assays were performed for each stimulation (∼40 cells/experiment).

**Figure EV4**. GCaMP6f responses of photoreceptor cells in isolated ommatidia before (basal) and after addition of 30 μM 2-linoleoyl glycerol (2-LG). The ommatidia were isolated from *ninaE*>*GCaMP6f* flies. Scale bars, 100 μm. **(A)** Control (*norpA*^*P24*^) ommatidia before addition of 2-LG. **(B)** Control ommatidia ∼240 seconds after addition of 2-LG. **(C)** *norpA*^*P24*^;*trpl*^*302*^;*trp*^*343*^ ommatidia before addition of 2-LG. **(D)** *norpA*^*P24*^;*trpl*^*302*^;*trp*^*343*^ ommatidia ∼240 seconds after addition of 2-LG.

## References

Abadji V, Lin S, Taha G, Griffin G, Stevenson LA, Pertwee RG, Makriyannis A (1994) (R)-methanandamide: a chiral novel anandamide possessing higher potency and metabolic stability. J Med Chem 37: 1889–1893

Acharya JK, Jalink K, Hardy RW, Hartenstein V, Zuker CS (1997) InsP_3_ receptor essential for growth and differentiation but not for vision in Drosophila. Neuron 18: 881–887

Asteriti S, Liu CH, Hardie RC (2017) Calcium signalling in Drosophila photoreceptors measured with GCaMP6f. Cell Calcium 65: 40–51

Bloomquist BT, Shortridge RD, Schneuwly S, Perdew M, Montell C, Steller H, Rubin G, Pak WL (1988) Isolation of a putative phospholipase C gene of Drosophila, norpA, and its role in phototransduction. Cell 54: 723–733

Bradshaw HB, Raboune S, Hollis JL (2013) Opportunistic activation of TRP receptors by endogenous lipids: exploiting lipidomics to understand TRP receptor cellular communication. Life Sci 92: 404–409

Bradshaw HB, Rimmerman N, Hu SS, Benton VM, Stuart JM, Masuda K, Cravatt BF, O’Dell DK, Walker JM (2009) The endocannabinoid anandamide is a precursor for the signaling lipid N-arachidonoyl glycine by two distinct pathways. BMC Biochem 10: 14

Britt SG, Feiler R, Kirschfeld K, Zuker CS (1993) Spectral tuning of rhodopsin and metarhodopsin in vivo. Neuron 11: 29–39

Chyb S, Raghu P, Hardie RC (1999) Polyunsaturated fatty acids activate the Drosophila lightsensitive channels TRP and TRPL. Nature 397: 255–259

Delgado R, Bacigalupo J (2009) Unitary recordings of TRP and TRPL channels from isolated Drosophila retinal photoreceptor rhabdomeres: activation by light and lipids. J Neurophysiol 101: 2372–2379

Delgado R, Delgado MG, Bastin-Héline L, Glavic A, O’Day PM, Bacigalupo J (2019) Light-induced opening of the TRP channel in isolated membrane patches excised from photosensitive microvilli from Drosophila photoreceptors. Neuroscience 396: 66–72

Delgado R, Muñoz Y, Peña-Cortés H, Giavalisco P, Bacigalupo J (2014) Diacylglycerol activates the light-dependent channel TRP in the photosensitive microvilli of Drosophila melanogaster photoreceptors. J Neurosci 34: 6679–6686

Elinder F, Liin SI (2017) Actions and mechanisms of polyunsaturated fatty acids on voltage-gated ion channels. Front Physiol 8: 43

Gregus AM, Buczynski MW (2020) Druggable targets in endocannabinoid signaling. Advances in experimental medicine and biology 1274: 177–201

Hardie RC (1991) Whole-cell recordings of the light-induced current in dissociated Drosophila photoreceptors - evidence for feedback by calcium permeating the lightsensitive channels. Proc Roy Soc Lond B Biol Sci 245: 203–210

Hardie RC, Franze K (2012) Photomechanical responses in Drosophila photoreceptors. Science 338: 260–263

Hardie RC, Juusola M (2015) Phototransduction in Drosophila. Curr Opin Neurobiol 34C: 37–45

Hardie RC, Minke B (1992) The trp gene is essential for a light-activated Ca^2^+ channel in Drosophila photoreceptors. Neuron 8: 643–651

Huang FD, Matthies HJ, Speese SD, Smith MA, Broadie K (2004) Rolling blackout, a newly identified PIP_2_-DAG pathway lipase required for Drosophila phototransduction. Nat Neurosci 7: 1070–1078

Huang J, Liu CH, Hughes SA, Postma M, Schwiening CJ, Hardie RC (2010) Activation of TRP channels by protons and phosphoinositide depletion in Drosophila photoreceptors. Curr Biol 20: 189–197

Inoue H, Yoshioka T, Hotta Y (1985) A genetic study of inositol trisphosphate involvement in phototransduction using Drosophila mutants. Biochem Biophys Res Commun 132: 513–519

Laine K, Jarvinen K, Mechoulam R, Breuer A, Jarvinen T (2002) Comparison of the enzymatic stability and intraocular pressure effects of 2-arachidonylglycerol and noladin ether, a novel putative endocannabinoid. Invest Ophthalmol Vis Sci 43: 3216–3222

Leung HT, Tseng-Crank J, Kim E, Mahapatra C, Shino S, Zhou Y, An L, Doerge RW, Pak WL (2008) DAG lipase activity is necessary for TRP channel regulation in Drosophila photoreceptors. Neuron 58: 884–896

Lev S, Katz B, Tzarfaty V, Minke B (2012) Signal-dependent hydrolysis of phosphatidylinositol 4,5-bisphosphate without activation of phospholipase C: Implications on gating of Drosophila TRPL (Transient Receptor Potential-Like) channel. J Biol Chem 287: 1436–1447

Lin G, Lee PT, Chen K, Mao D, Tan KL, Zuo Z, Lin WW, Wang L, Bellen HJ (2018) Phospholipase PLA2G6, a Parkinsonism-associated gene, affects Vps26 and Vps35, retromer function, and ceramide levels, similar to α-synuclein gain Cell Metab 28: 605–618 e606

Liu J, Wang L, Harvey-White J, Osei-Hyiaman D, Razdan R, Gong Q, Chan AC, Zhou Z, Huang BX, Kim HY et al (2006) A biosynthetic pathway for anandamide. Proc Natl Acad Sci USA 103: 13345–13350

Liu L, MacKenzie KR, Putluri N, Maletic-Savatic M, Bellen HJ (2017) The glia-neuron lactate shuttle and elevated ROS promote lipid synthesis in neurons and lipid droplet accumulation in glia via APOE/D. Cell Metab 26: 719–737 e716

Long JZ, Li W, Booker L, Burston JJ, Kinsey SG, Schlosburg JE, Pavon FJ, Serrano AM, Selley DE, Parsons LH et al (2009) Selective blockade of 2-arachidonoylglycerol hydrolysis produces cannabinoid behavioral effects. Nat Chem Biol 5: 37–44

Marza E, Long T, Saiardi A, Sumakovic M, Eimer S, Hall DH, Lesa GM (2008) Polyunsaturated fatty acids influence synaptojanin localization to regulate synaptic vesicle recycling. Mol Biol Cell 19: 833–842

McGurk L, Berson A, Bonini NM (2015) Drosophila as an in vivo model for human neurodegenerative disease. Genetics 201: 377–402

McPartland J, Di Marzo V, De Petrocellis L, Mercer A, Glass M (2001) Cannabinoid receptors are absent in insects. J Comp Neurol 436: 423–429

Minke B (1982) Light-induced reduction in excitation efficiency in the trp mutant of Drosophila. J Gen Physiol 79: 361–385

Minke B, Wu C, Pak WL (1975) Induction of photoreceptor voltage noise in the dark in Drosophila mutant. Nature 258: 84–87

Montell C (2012) Drosophila visual transduction. Trends Neurosci 35: 356–363

Montell C (2021) Drosophila sensory receptors—a set of molecular Swiss Army Knives. Genetics 217: 1–34

Montell C, Rubin GM (1989) Molecular characterization of the Drosophila trp locus: a putative integral membrane protein required for phototransduction. Neuron 2: 1313–1323

Muller C, Lynch DL, Hurst DP, Reggio PH (2020) A closer look at anandamide interaction with TRPV1. Front Mol Biosci 7: 144

Muller C, Morales P, Reggio PH (2019) Cannabinoid ligands targeting TRP channels. Front Mol Neurosci 11: 487

Muller C, Reggio PH (2020) An analysis of the putative CBD binding site in the ionotropic cannabinoid receptors. Front Cell Neurosci 14: 615811

Niemeyer BA, Suzuki E, Scott K, Jalink K, Zuker CS (1996) The Drosophila lightactivated conductance is composed of the two channels TRP and TRPL. Cell 85: 651–659

Nomura DK, Blankman JL, Simon GM, Fujioka K, Issa RS, Ward AM, Cravatt BF, Casida JE (2008) Activation of the endocannabinoid system by organophosphorus nerve agents. Nat Chem Biol 4: 373–378

Pak WL (1994) Retinal degeneration mutants of Drosophila. In: Molecular genetics of inherited eye disorders, Wright A.F., Jay B. (eds.) pp. 29–52. Harwood Academic Publishers: Chur, Switzerland

Pak WL, Grossfield J, Arnold KS (1970) Mutants of the visual pathway of Drosophila melanogaster. Nature 227: 518–520

Parnas M, Katz B, Minke B (2007) Open channel block by Ca^2^+ underlies the voltage dependence of Drosophila TRPL channel. J Gen Physiol 129: 17–28

Phillips AM, Bull A, Kelly LE (1992) Identification of a Drosophila gene encoding a calmodulin-binding protein with homology to the trp phototransduction gene. Neuron 8: 631–642

Pumroy RA, Samanta A, Liu Y, Hughes TE, Zhao S, Yudin Y, Rohacs T, Han S, Moiseenkova-Bell VY (2019) Molecular mechanism of TRPV2 channel modulation by cannabidiol. eLife 8

Raghu P, Hardie RC (2009) Regulation of Drosophila TRPC channels by lipid messengers. Cell Calcium 45: 566–573

Raghu P, Usher K, Jonas S, Chyb S, Polyanovsky A, Hardie RC (2000) Constitutive activity of the light-sensitive channels TRP and TRPL in the Drosophila diacylglycerol kinase mutant, rdgA. Neuron 26: 169–179

Ranganathan R, Harris GL, Stevens CF, Zuker CS (1991) A Drosophila mutant defective in extracellular calcium-dependent photoreceptor deactivation and desensitization. Nature 354: 230–232

Schindelin J, Arganda-Carreras I, Frise E, Kaynig V, Longair M, Pietzsch T, Preibisch S, Rueden C, Saalfeld S, Schmid B et al (2012) Fiji: an open-source platform for biological-image analysis. Nat Methods 9: 676–682

Shen LR, Lai CQ, Feng X, Parnell LD, Wan JB, Wang JD, Li D, Ordovas JM, Kang JX (2010) Drosophila lacks C20 and C22 PUFAs. J Lipid Res 51: 2985–2992

Tortoriello G, Beiersdorf J, Romani S, Williams G, Cameron GA, Mackie K, Williams MJ, Di Marzo V, Keimpema E, Doherty P et al (2021) Genetic manipulation of sn-1-diacylglycerol lipase and CB_1_ cannabinoid receptor gain-of-function uncover neuronal 2-linoleoyl glycerol signaling in Drosophila melanogaster. Cannabis Cannabinoid Res 6: 119–136.

Tortoriello G, Rhodes BP, Takacs SM, Stuart JM, Basnet A, Raboune S, Widlanski TS, Doherty P, Harkany T, Bradshaw HB (2013) Targeted lipidomics in Drosophila melanogaster identifies novel 2-monoacylglycerols and N-acyl amides. PLoS One 8: e67865

Wang T, Jiao Y, Montell C (2005a) Dissecting independent channel and scaffolding roles of the Drosophila transient receptor potential channel. J Cell Biol 171: 685–694

Wang T, Xu H, Oberwinkler J, Gu Y, Hardie RC, Montell C (2005b) Light activation, adaptation, and cell survival functions of the Na+/Ca^2^+ exchanger CalX. Neuron 45: 367–378

Warrick JM, Paulson HL, Gray-Board GL, Bui QT, Fischbeck KH, Pittman RN, Bonini NM (1998) Expanded polyglutamine protein forms nuclear inclusions and causes neural degeneration in Drosophila. Cell 93: 939–949

Wes PD, Chevesich J, Jeromin A, Rosenberg C, Stetten G, Montell C (1995) TRPC1, a human homolog of a Drosophila store-operated channel. Proc Natl Acad Sci USA 92: 9652–9656

Yau KW, Hardie RC (2009) Phototransduction motifs and variations. Cell 139: 246-264

Yoshioka T, Inoue H, Kasama T, Seyama Y, Nakashima S, Nozawa Y, Hotta Y (1985) Evidence that arachidonic acid is deficient in phosphatidylinositol of Drosophila heads. J Biochem 98: 657–662

Zhu X, Chu PB, Peyton M, Birnbaumer L (1995) Molecular cloning of a widely expressed human homologue for the Drosophila trp gene. FEBS Lett 373: 193–198

Zhuang N, Li L, Chen S, Wang T (2016) PINK1-dependent phosphorylation of PINK1 and Parkin is essential for mitochondrial quality control. Cell Death Dis 7: e2501

Ziegler AB, Menage C, Gregoire S, Garcia T, Ferveur JF, Bretillon L, Grosjean Y (2015) Lack of dietary polyunsaturated fatty acids causes synapse dysfunction in the Drosophila visual system. PLoS One 10: e0135353

