## Supplementary figures and images for "Light-induction of endocannabinoids and activation of *Drosophila* TRPC channels"

### Supplemental Figure 1

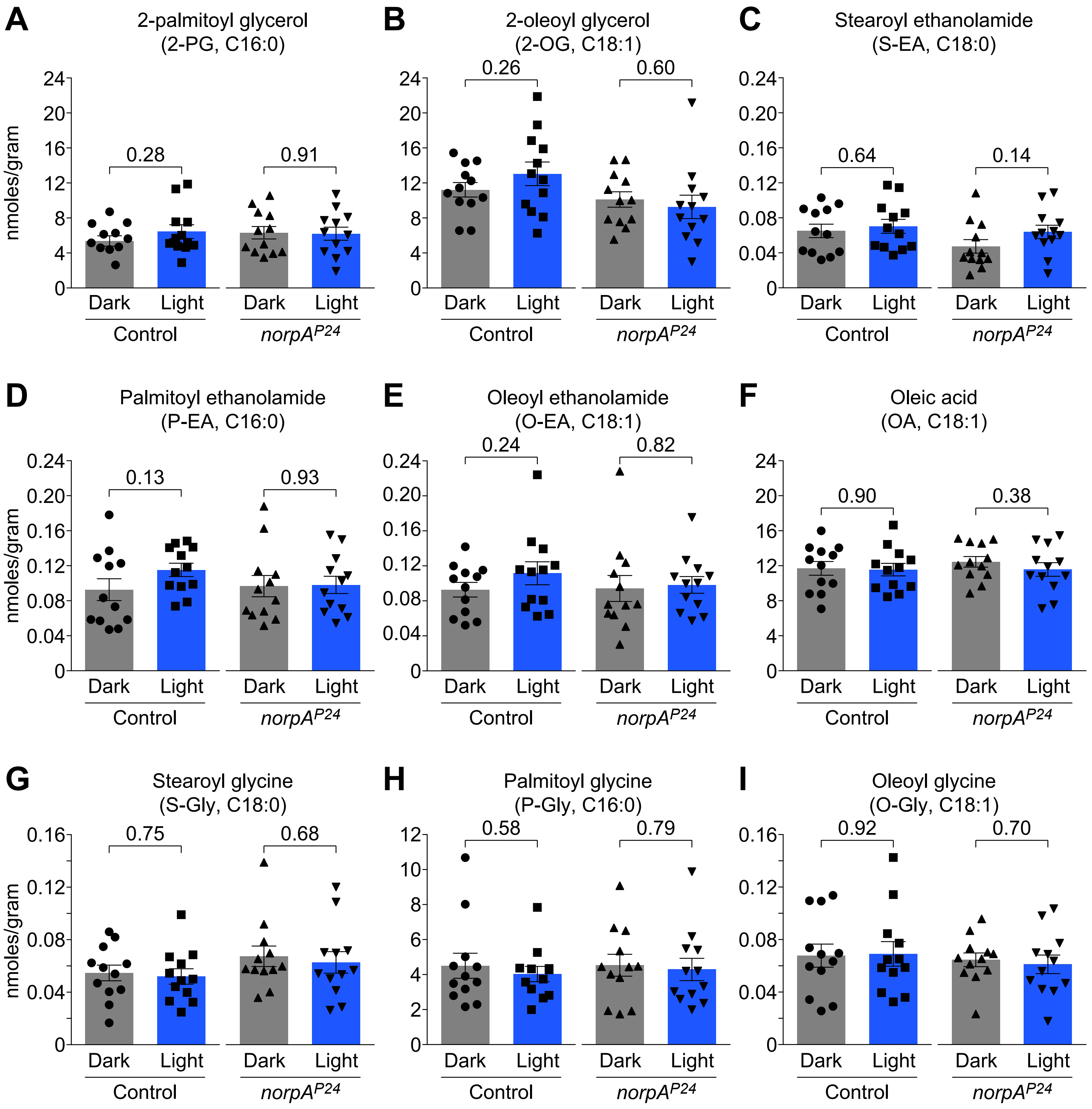

### Supplemental Figure 2

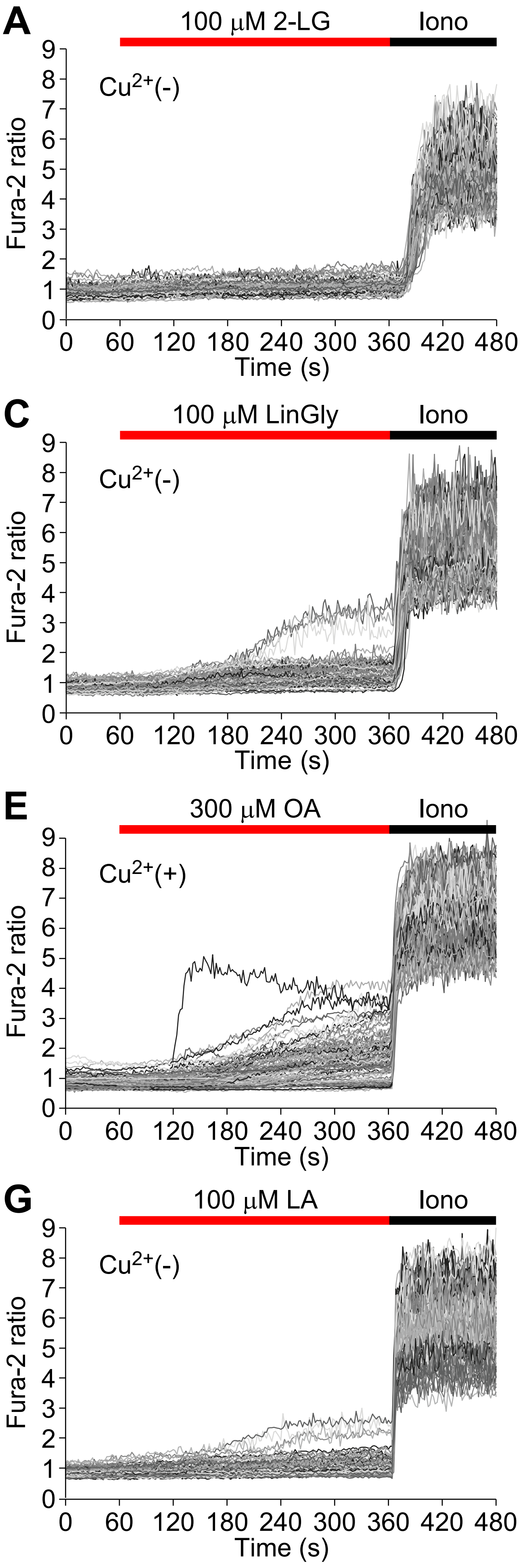

### Supplemental Figure 3

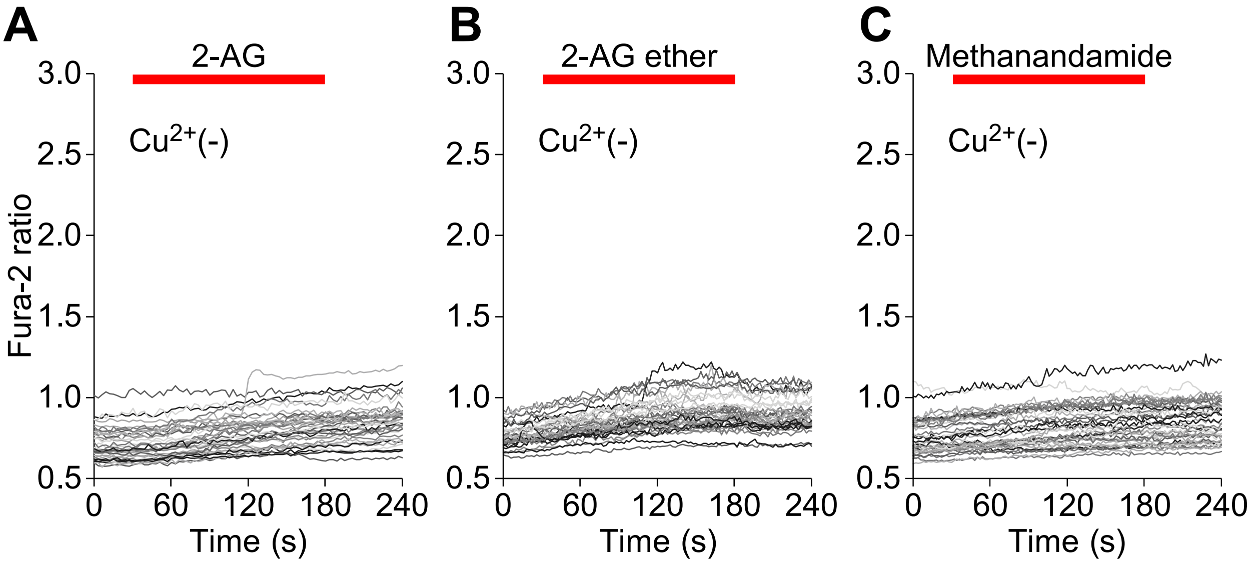

### Supplemental Figure 4

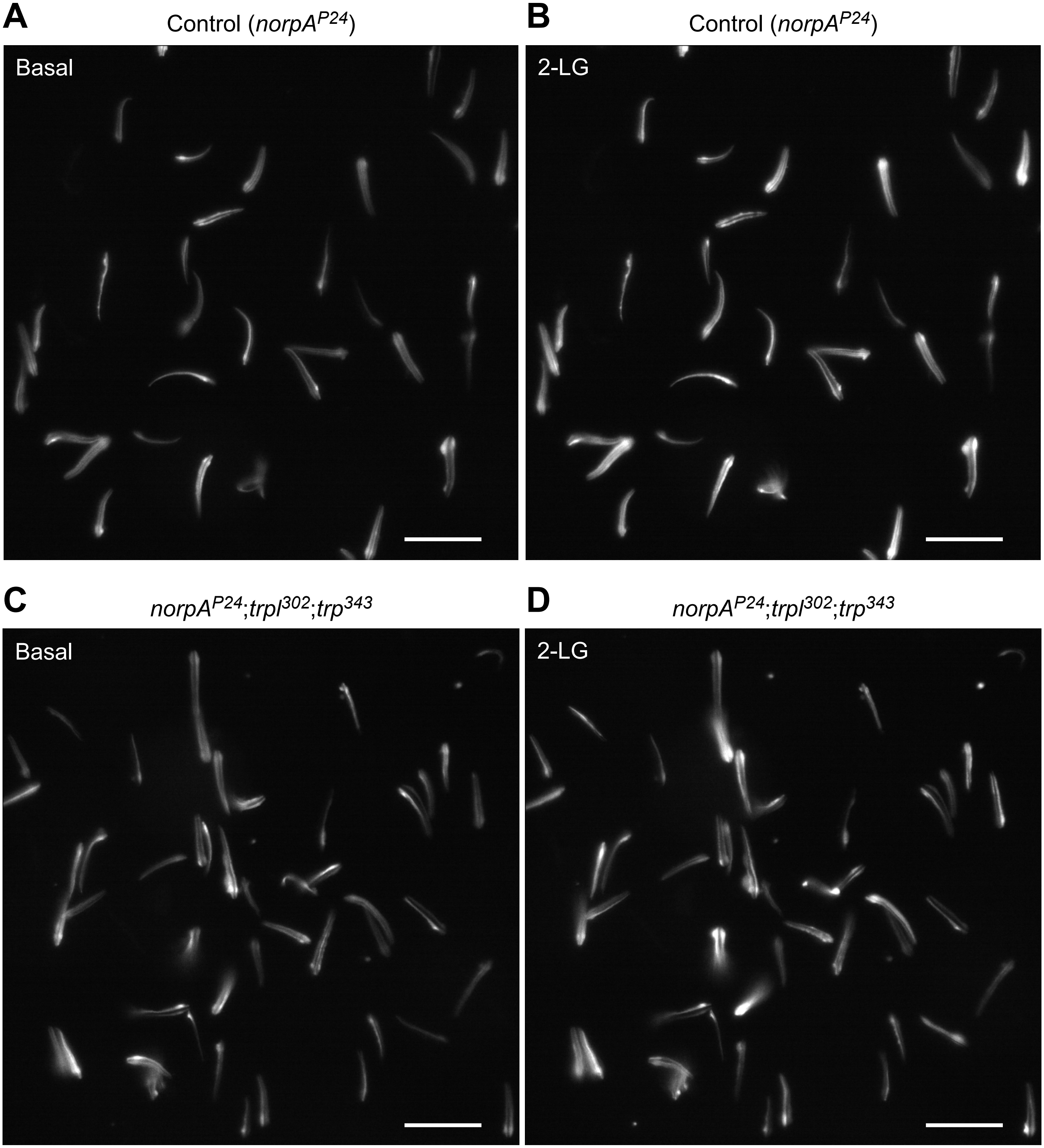
